# CS-Fold: Advancing RNA Structure Predictions through Phylogenetic Modelling of Compensatory Mutations in Deep Neural Networks

**DOI:** 10.1101/2025.04.27.650904

**Authors:** Jiren Zhou, Jiajia Xu, Jiayu Wen, Brian John Parker

## Abstract

Accurate prediction of RNA secondary structures is essential for understanding the conformation, function, and interactions of RNA. Leveraging co-evolutionary information across species through multiple sequence alignments (MSAs) has been proven to be effective in improving molecular structure predictions. However, existing deep learning approaches do not explicitly incorporate compensatory substitutions along the phylogenetic trees, which are crucial for capturing structural conservation through evolution. To address this, we developed **CS-Fold**, a novel deep learning framework that integrates compensatory substitutions likelihoods as constraints using likelihood estimation and Monte Carlo algorithms. These likelihoods, representing evolutionary changes along phylogenetic trees, are encoded in a sparse matrix that guides the attention mechanism of a Pairformer-based architecture. A custom loss function and an unrolled post-processing algorithm enforce adherence to the solution space constrained by these evolutionary constraints. **CS-Fold** achieves a substantial 5% improvement in the F1 score compared to the current mainstream approaches, demonstrated through evaluations on cross-family datasets, including 604 human RNA families from the Rfam database. Our model offers novel insights and incorporates additional evolutionary information beyond traditional MSAs and folding strategies, providing a robust and innovative solution for RNA secondary structure prediction.

## 1 Introduction

RNA secondary structure refers to the folded structures formed by a single RNA strand through complementary base pairing. These structures are crucial role for regulating RNA stability, activity, and molecular interactions of RNAs [1]. While experimental techniques such as Nuclear magnetic resonance (NMR) and X-ray crystallography can be used to analyze RNA structures at the biochemical level [2], they are costly and resource-intensive. Consequently, computational approaches for RNA structure prediction are essential. In RNA structure modeling, secondary structure reveals the fundamental nested base-pairing interactions that underpin RNA function, whereas tertiary structure is important for understanding RNA-protein interactions. Given that RNA secondary structure is more strongly constrained by base pairing than its tertiary structure, we focus on accurately predicting stable RNA secondary structures to provide a foundation for future analyses of complex RNA interactions.

A range of deep learning methods have been applied to RNA secondary structure prediction. Among them, MXFold2 [3] leverages the Minimum Free Energy (MFE) hypothesis, incorporating Turner’s empirical energy parameters [4]. By using deep neural networks, MXFold2 enhances the prediction of dynamic programming-based structure, improving the accuracy and efficiency of the RNA folding analysis. E2Efold [5] employs an unrolled optimization algorithm to incorporate constraints such as minimum loop size, illegal pairings, and non-repetitive pairings as output constraints of the deep network, effectively reducing the solution space. Additionally, architectures that rely on supervised learning, such as UFold [6], transform RNA secondary structure prediction into an image-based problem and adapt the U-Net architecture to perform predictions, effectively framing it as an image segmentation task. Recently, advancements in Large Language Model (LLM) have led to the development of pretrained sequence models that include RNA secondary structure prediction as one of their downstream tasks. Models such as ERNIE-RNA [7], RNAErnie [8], and RiNALMo [9] leveraging the scaling capabilities of LLMs to serve as powerful semantic feature extractors, enhancing RNA structure prediction accuracy. However, existing methods still lack a framework that explicitly incorporates compensatory substitution information—an important co-evolutionary signal unique to RNA that distinguishes from protein folding. Incorporating this biological prior into deep learning presents a critical opportunity to improve prediction performance.

As shown in Figure 1, existing methods for RNA secondary structure prediction rely on unsupervised learning from column-wise frequency in MSAs. However, simply using MSA is limited, as it does not account for phylogenetic tree topology and branch lengths, making it suboptimal for capturing evolutionary constraints in RNA. Structured RNAs evolve through characteristic substitution patterns that preserve base-pair interactions. These include compensatory double substitutions which maintain the Watson-Crick G:C and A:U base pairs (e.g., *A* : *U → G* : *C*), and compatible single substitutions which maintain G:U wobble base pairs by a single mutation (e.g., *U* : *G → U* : *A*): these maintain the stability of the RNA structure, while incompatible mutations degrade it. Extracting these compensatory signals is crucial but challenging due to their sparsity and the limitations of traditional MSA-based models.

**Figure 1:**
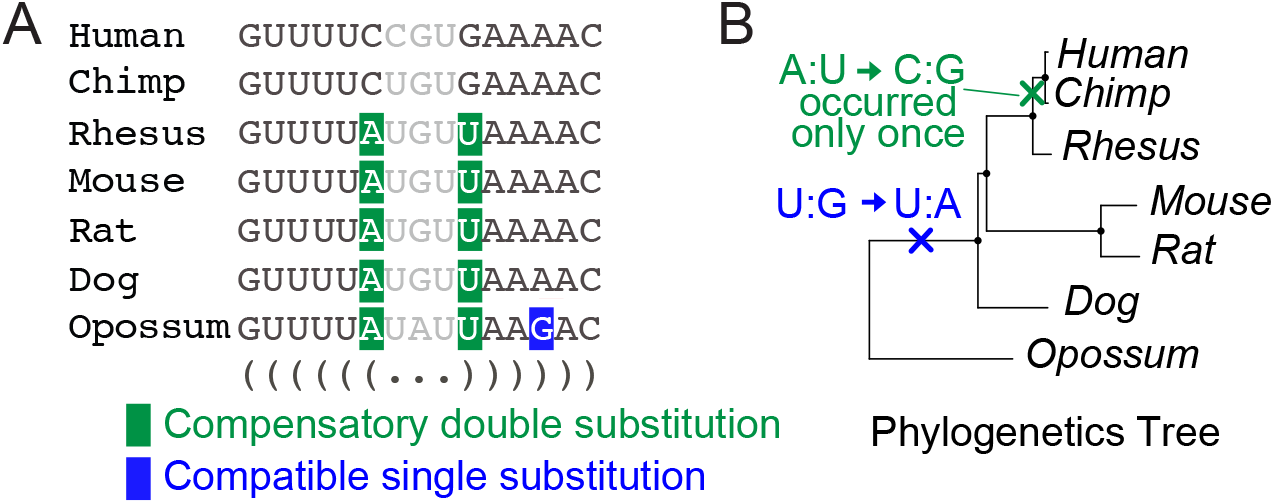
Mutation evidence cannot be quantified by MSA alone. **A**. An example MSA highlights compensatory double substitutions (green) and compatible single substitutions (blue), with corresponding structure in dot-bracket notation. **B**. Corresponding phylogenetic tree shows that the most parsimonious explanation for the mutations in MSA A is a single A:U to C:G compensatory double substitution after the last common ancestor of human and chimp, rather than five independent mutations in Rhesus, Mouse, Rat, Dog, Opossum as suggested by the MSA. This highlights the essential need for full phylogenetic modelling of the mutation evidence.

Our approach addresses this limitation by reconstructing the phylogenetic ancestral sequences within the phylogenetic tree to extract compensatory substitutions—sparse but biologically significant signals that are obscured in traditional MSA-based methods. By leveraging a Monte Carlo algorithm over the phylogenetic tree, we model these evolutionary signals more correctly. Specifically, we calculate p-values, *p* for both compensatory double and single compatible single substitutions at each site during evolution and use them as constraints to preserve the phylogenetic tree’s topological information of RNAs. This explicit incorporation of evolutionary structure reduces the search space for deep learning models, making their outputs more biologically grounded and computationally efficient.

We define our problem as solving a symmetric duality relationship in RNA secondary structure prediction. To address this, we designed a deep learning framework based on triangular updates, incorporating triangular multiplication and triangular attention as its core components. Given the sparsity of compensatory substitution signals, we introduced attention bias to two of the four heads in triangular multi-head attention. This guides multi-head symmetric attention to update from two complementary perspectives: one capturing compensatory substitution signals, and the other leveraging a global receptive field.

Additionally, we designed a custom loss function and applied an optimization-based approach during inference. Specifically, we imposed a direct gradient penalty on model outputs for sites exhibiting unilateral or bilateral compensatory substitutions, ensuring that outputs fall below 1 ™ *p*. This end-to-end constraint effectively refines the model’s prediction output within a biologically meaningful solution space.

We summarize our contributions to this field:

- We utilized the vertebrate evolutionary tree topology to reconstruct ancestral sequences using a maximum likelihood method. Through Monte Carlo simulation of the evolutionary process, p-values were calculated for both compatible and compensatory double and single mutations at each base pair. This approach addresses the absence of phylogenetic tree structure in existing MSA-based algorithms and leverages 1 − *p* as a calibrated prior for the deep learning framework.
- We formulated the problem as a symmetric duality relationship problem and developed a deep learning framework that incorporates triangular multiplication and triangular attention to explicitly model this duality relationships. This design provides an attention-based prior for deep learning, establishing a paradigm that can be extended to other biomolecular structure prediction tasks.
- We applied optimization-based constraints to ensure that model outputs at specific sites remain below the prior knowledge threshold defined by piror knowledge. This constraint constrains effectively narrows the solution space during inference, leading to more biologically meaningful and robust predictions.

## 2 Related Work

### 2.1 Method for reconstructing the phylogenetic ancestral sequences of RNAs

PAML (Phylogenetic Analysis by Maximum Likelihood) [10] is a widely used method for reconstructing ancestral sequences in phylogenetic trees using maximum likelihood estimation. In our study, we employ PAML to reconstruct ancestral RNA sequences using a 100-way vertebrate species phylogenetic tree and multiple sequence alignments obtained inferred from human RNA family sequences in Rfam. Leveraging the reconstructed sequences and tree structure, we perform statistical analysis of compensatory substitutions across all possible base pairings through Monte Carlo evolutionary modeling.

### 2.2 Dual Representation Update

The problem was defined as a sparse symmetric duality problem, where each site’s embedding represents its interactions with other sites. The Pairformer mechanism, originally used in AlphaFold 2 and 3 [11, 12], which was designed for preserving structural consistency, was employed for update embedding. The triangular attention block incorporates symmetric attention along with explicit attention biasing. During incoming and outgoing updates, we apply a transposed prior, ensuring that prior information is effectively utilized in both directions to enhance model expressiveness and stability.

### 2.3 Unrolled Optimization

During inference, we apply an unrolled algorithm-based post-processing step to further constrain the model’s output. Building on E2Efold’s [5] constraints—enforcing legal base-pairing, minimum loop size, and single RNA pairings—we introduce an additional penalty mechanism. Specifically, for model outputs where the predicted probability falls below 1 - p, we apply gradient penalties to guide predictions toward compliance with biological hard constraints and compensatory mutation priors. Our design also references other differential methods for RNA structure prediction [13, 14].

## 3 Compensatory Substitutions Statistics

First, define the human reference genome (assembly hg38) with MSA information as ℛ_hg38_, for ∀*r* ∈ ℛ_hg38_, *r* represents the reference human rna, there exists a set ℳ_*r*_ ⊆ 𝒮_100_.

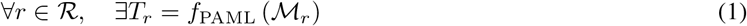

*f*_PAML_ - PAML function to reconstruct the phylogentics tree ancestral sequences.

*T*_*r*_ - The phylogenetic tree reconstructed for each reference RNA, exists *T*_*r*_ = (*V, E*). *V* define as set of node. *E* define as set of edge.

Here, we obtain 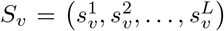, which represents the set of sequences inferred at all nodes, where *L* is the length of *r*.

For any *s* in *S*_*v*_, any position *s*_*i*_, *i* ∈ {1, 2, …, *L*}

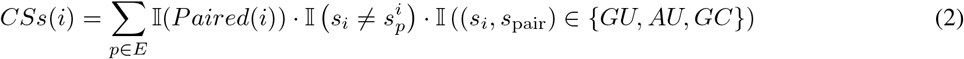

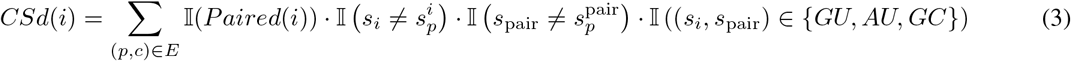

Here, 𝕀(*Paired*(*i*)) indicates that the base must be in a paired region. When the corresponding position in the child node differs from the parent node but still maintains pairing with its counterpart, it is defined as single substitution. It follows that if the paired counterpart also differs from the parent node while still maintaining a valid pairing, it constitutes double substitution.

*s*_anc_ is the inferred ancestor sequence for *T*_*r*_,

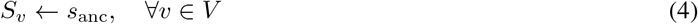

Set the sequences of all nodes in the tree to the inferred ancestral sequences.

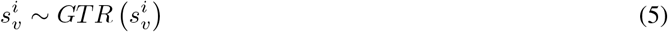

Apply GTR to generate the same number of mutations as before at all positions of all sequences. For any *k*_th_ iteration, the following observation exists.

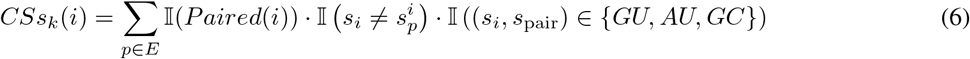

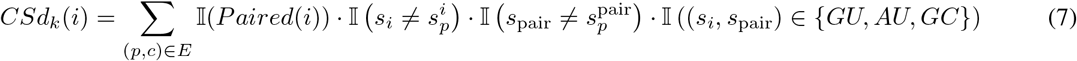

For any *r*, the following two statistical values exist:

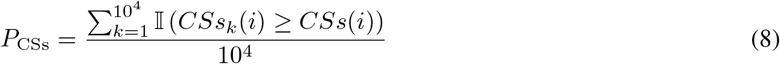

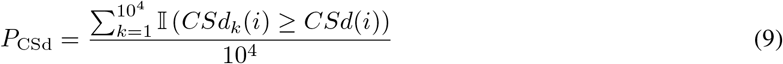

## 4 Pairformer

As defined in the previous section, the final *A* should be sparse and symmetric. We introduce the Pairformer architecture specifically designed to handle dual problems. Each module consists of outgoing triangle update, incoming triangle update, triangle self-attention around the starting node, triangle self-attention around the ending node, and a final transition step. We stack two layers of this structure.

The remaining parts of the model mainly consist of an MLP for dimensionality reduction and a pre-trained large model that provides the initial embeddings for the architecture. Here, we use RiNALMo [9], which has not been fine-tuned for RNA secondary structure prediction, and we freeze all weights, using only its inference results.

Specifically, as shown in Figure 2, we design the attention mechanism as follows:

**Figure 2:**
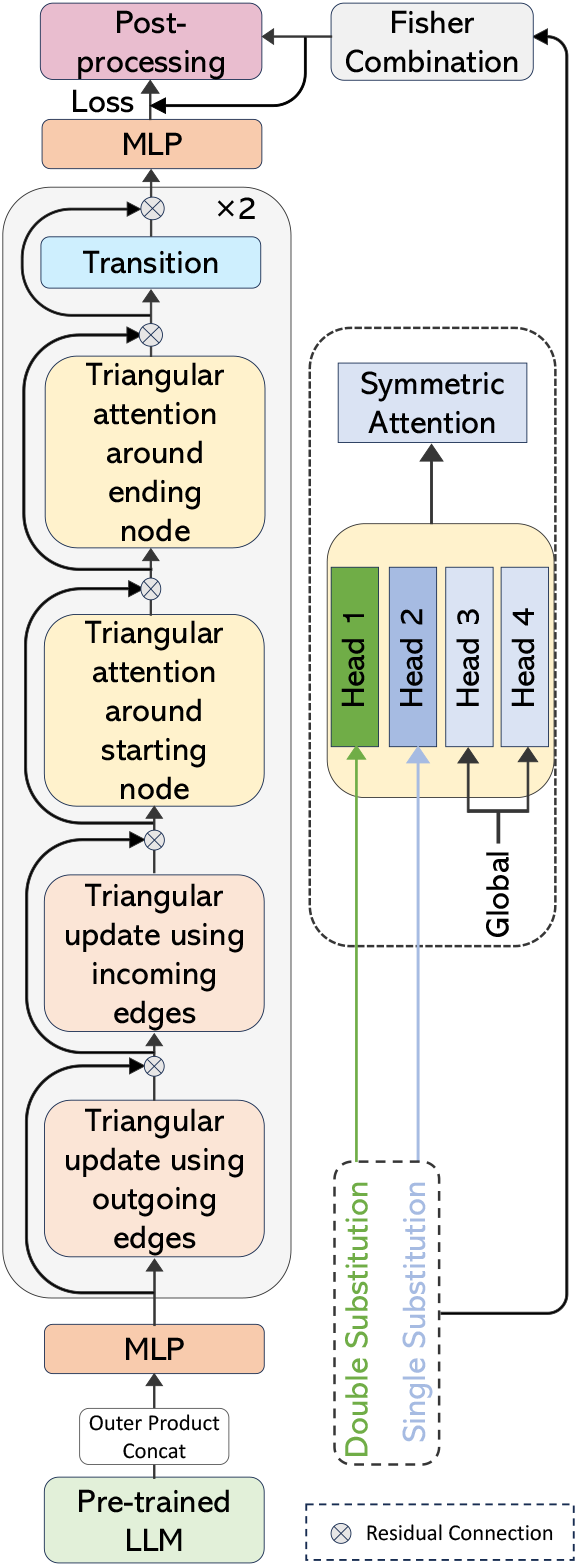
The architecture diagram of the CS-Fold model.

**Figure 3:**
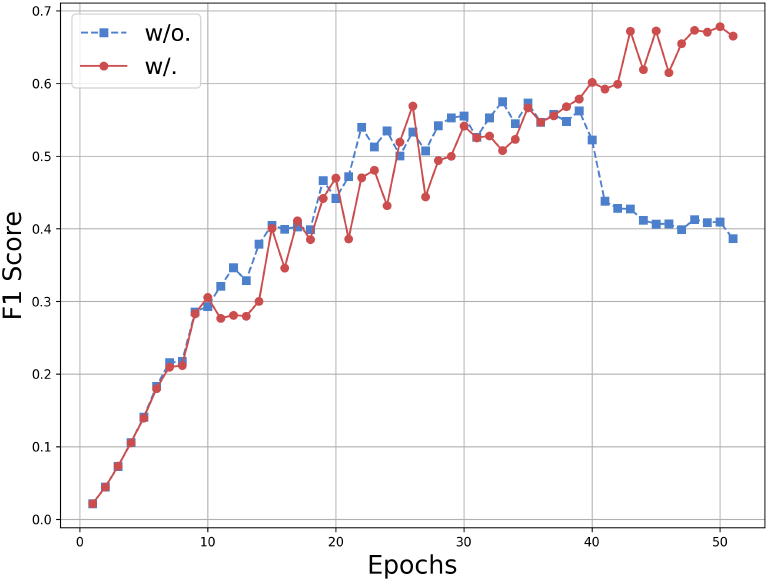
The comparison of the first 51 epochs between with phylogenetic information and without

- The attention remains symmetric during updates.

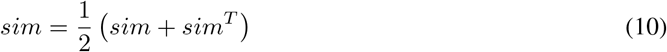
- We use 4 attention heads. The first two heads incorporate prior knowledge obtained earlier as attention biases, guiding the network to focus on these special sites. The remaining two heads maintain a global receptive field to capture broader information. Since this information is extremely sparse, it serves only as a guiding signal rather than a definitive solution.

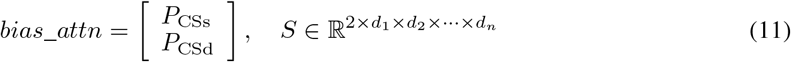

## 5 Post Processing

We first apply Fisher’s combined test to *P*_CSs_ and *P*_CSd_ to merge them into a variable in the post-processing step.

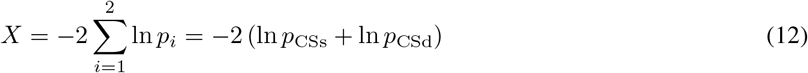

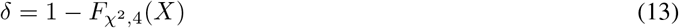

First, we define the optimization objective and the constraints that need to be satisfied.

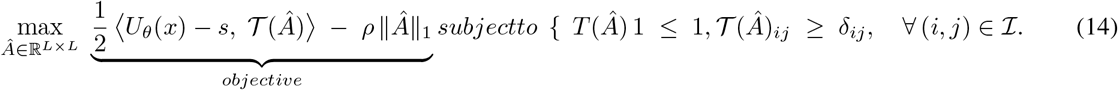

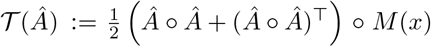, where follows e2efold for transformation. Matrix *M* is defined as ℐ represents the set that only includes positions with compensatory substitutions. This is crucial because these values cannot be set to 0. 1 − *p* indicates non-significance; however, in this context, we need to define meaningless on most positions. To handle these constraints, two sets of dual variables were given, 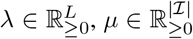. the Lagrangian ℒ could be defined as below:

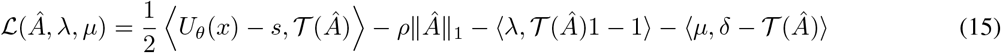

Here, use the primal-dual solution.

### Primal gradient step

Update *Â* in the direction of the (negative) gradient of the Lagrangian with respect to *Â*. Let *α* be the primal step size, and *γ*_*α*_ a decay factor. Then

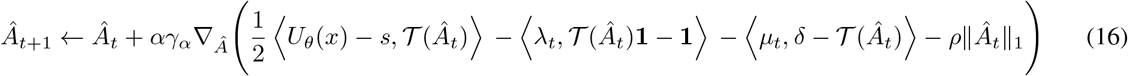

The *𝓁*_1_-term ∥*Â*_*t*_∥_1_ is not differentiable at 0, so we use a soft-threshold.

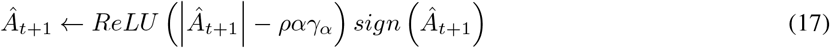

Then map back to *A*.

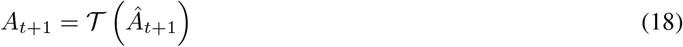

### Dual gradient update for *λ*

*λ* could be updated by subgradient ascent on the dual objective. If we denote *β* as the dual step size and *γ*_*β*_ as its decay, then

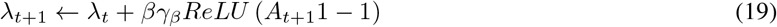

### Dual gradient update for *µ*

*µ* could be updated by subgradient ascent on the dual objective. If we denote *β* as the dual step size and *γ*_*β*_ as its decay, then

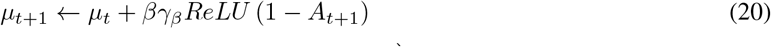

Intuitively, if *A*_*ij*_ in iteration *t* + 1 violates the threshold *δ*_*ij*_ (i.e., 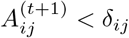), then *µ*_*ij*_ gets pushed up in the next primal gradient step.

## 6 Loss Function

Here, we use a composite loss function to train the model:

- Weighted BCE (Binary Cross-Entropy): We assign a higher weight to the positive class.
- Tversky Loss: This loss is used to balance precision and recall. During training, we observed that the model initially tends to produce an excessively high recall, leading to nearly all outputs being classified as positive.
- Constraint-based loss on specific positions: This ensures that the model adheres to the required constraints at the positional level.

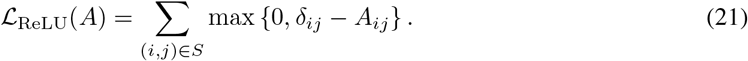

Overall loss function is as shown below:

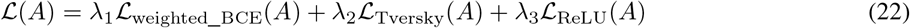

## 7 Experiment

### 7.1 Dataset

Current approaches for RNA secondary structure prediction rely heavily on datasets like ArchiveII [15] and TS0 [16] within RNA families, and PDB [17] and bpRNA_new [18] across families. However, these datasets are predominantly derived from bacterial RNAs, limiting their utility for comprehensive analysis of RNA families in other organisms. Bacterial phylogenies, shaped by rapid evolution and horizontal gene transfer, lack the deep, hierarchical branching patterns found in vertebrate phylogenies, complicating evolutionary comparisons.

To overcome this limitation and advance the study of vertebrate RNA families, we created a new dataset based on vertebrate RNA families. We obtained human RNA family sequences with their structures from Rfam 14.5, along with the 100-way vertebrate species tree and multiple sequence alignments from UCSC. Using mafFrag, we extracted a total of 13,962 vertebrate RNA sequences and reconstructed them with PAML, successfully inferring 13,898 sequences. A small number of RNAs could not be reconstructed due to an insufficient homologous sequences in the phylogenetic tree. To accommodate the processing limitations of pre-trained models, sequences longer than 1,028 nucleotides were excluded, yieldinga final dataset of 13,778 sequences. These sequences span 604 RNA families, with 9 families showing disproportionately large representation. The Metazoa_SRP family was the most abundant, comprising 5,496 sequences. Detailed statistics for these 9 families are provided in Table 1.

**Table 1:** Dominant RNA family in our Datasets.

| FAMILY | NUMBER | PROPORTION |
| --- | --- | --- |
| METAZOA_SRP | 5496 | 39.9 |
| Y_RNA | 744 | 5.4 |
| U6 | 1244 | 9.0 |
| TRNA | 696 | 5.1 |
| MIR1255 | 1220 | 8.8 |
| 7SK | 300 | 2.0 |
| MIR548 | 211 | 1.5 |
| 5S_rRNA | 390 | 2.8 |
| U7 | 209 | 1.5 |
| **TOTAL** | **10510** | **76.3** |

### 7.2 Data Augmentation

To enhance the diversity of training samples and mitigate overfitting, we employed two data augmentation strategies:

#### Downsampling

For 9 families with more than 200 sequences, 5 families were randomly assigned to the validation and test sets, while the remaining 4 families were included in the training set. We ensured no overlap between sets. All 9 families were downsampled to 200 sequences each after selection.

#### Geometric Augmentation

In the training set, we applied mirroring to both sequences and structures. This transformation preserved RNA structural integrity while altering sequential semantics, thereby affecting nucleotide positioning for pre-trained models.

In the training set, we applied mirroring to both sequences and structures. This transformation preserved RNA structural integrity while altering sequential semantics, thereby affecting nucleotide positioning for pre-trained models.

### 7.3 Metrics

We define the evaluation metrics at the per-base level as follows:

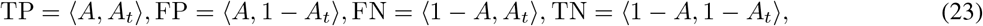

where *A* denotes the predicted base pairing, and *A*_*t*_ represents the true base pairing label. The true positive (TP), false positive (FP), false negative (FN), and true negative (TN) values are computed for each base in the RNA structure.

### 7.4 Model Performance

We compared our approach with several mainstream deep learning methods for RNA secondary structure prediction. These included MXfold2, which integrates deep networks with minimum free energy scoring; E2Efold, a transformer-based model with biological constraints that influenced our post-processing strategy; UFold, which uses image-based representations; and Rinalmo and ERNIE-RNA, both fine-tuned large language model (LLM) approaches.

Our model outperformed the current state-of-the-art, Rinalmo, by achieving a 5% higher F1 score (Table 2). Notably, unlike LLM-based models that require extensive fine-tuning, our method relies on a semantic masking strategy without depending on pre-trained models. In our evaluation, E2Efold and UFold were re-trained on our dataset, while MXfold2 was assessed using its pre-trained model. As shown in Table 2, our model demonstrates superior performance in precision and F1-score, though it does not show an advantage in recall. Precision is particularly critical for this task, where accurately identifying key structural elements is more important than maximizing overall recall. We excluded E2Efold’s results from the final comparison due to its known generalization issues with small cross-family datasets. These limitations are inherent to transformer-based networks, which require large amounts of training data. However, E2Efold’s post-processing remains valuable for constraining biologically unrealistic predictions.

**Table 2:** RNA structure prediction results.

| FAMILY | PRECISION | RECALL | F1 |
| --- | --- | --- | --- |
| **CSFOLD** | **78.8** | 77.0 | **77.9** |
| UFOLD | 57.7 | **83.0** | 65.5 |
| MXFOLD2 | 62.2 | 73.4 | 67.3 |
| RINALMO | 67.4 | 82.5 | 73.3 |
| ERNIE-RNA | 58.3 | 69.2 | 63.3 |
| E2EFOLD | - | - | - |

To mitigate data dependency issues common to attention-based models, we designed a lightweight neural network architecture with several optimizations:

- **Memory and Receptive Field Optimization:**. To balance memory capacity while maintaining a global receptive field without using windowed attention, we reduced the hidden dimensionality to 64 after the outer product concatenation for Pairformer updates. We stacked only two layers of Pairformer, compared to AlphaFold’s 48-layer network with a recycling mechanism during inference.
- **Semantic Feature Extraction:**Rinalmo, a 6.5B model that was not fine-tuned specifically for secondary structure prediction, was employed as our semantic extractor. We also compared this model after fine-tuning, demonstrating that its inference capability was not the key factor contributing to the strong performance of our model.

### 7.5 sAblation

Since our model integrates biases in attention, loss design, and post-processing constraints simultaneously, we conducted an ablation study to assess the impact of excluding compensatory substitution priors at all phases. The comparison results are presented in Table 3.

**Table 3:**
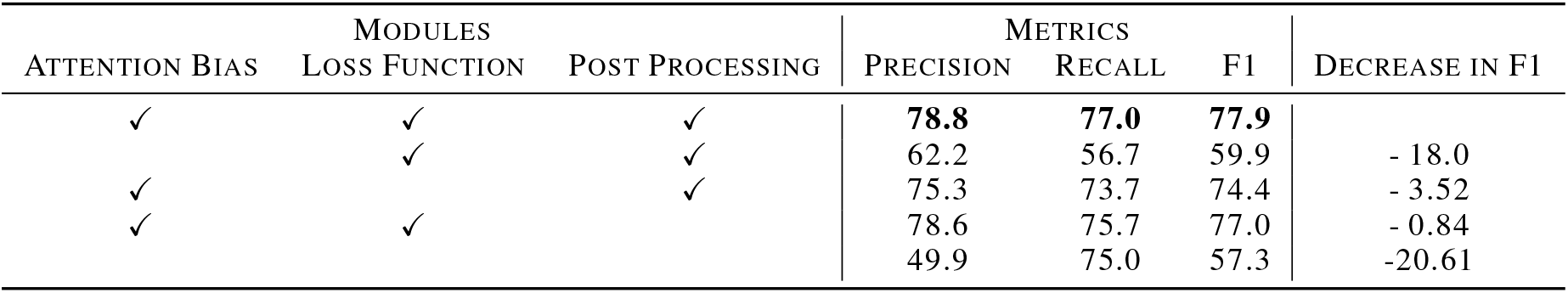
Ablation study with and without phylogenetic information.

The results show a significant performance drop when this prior information is omitted. We measured this effect using the F1 score on the validation set across training epochs. Without the compensatory substitution prior, the network peaked in performance around epoch 35 before gradually overfitting. In contrast, when the prior was included, the network exhibited more frequent performance spikes. This behavior is likely due to the smaller learning rate we applied, allowing the model to gradually escape local minima and improve its overall performance over time.

### 7.6 Data and source code availability

The data and source code will be made publicly available upon acceptance of the manuscript.

## 8 Conclusion

While MSA-based methods have improved RNA secondary structure prediction, they do not incorporate a sophisticated evolutionary model, and instead are based on frequency statistics within aligned sequence columns. However, this approach inherently leads to the loss of phylogenetic tree topology information, as bases with the same frequency in a column may have emerged at completely different evolutionary stages during co-evolution.

Our method, **CS-Fold**, overcomes this limitation by extracting and preserving phylogenetic information from MSA. We achieve this through a dually-updated network architecture with targeted loss functions and post-processing constraints that impose both soft and hard limits, maintaining evolutionary context throughout the prediction process.

Our experimental results demonstrate that integrating biological prior knowledge into deep learning models for RNA secondary structure prediction not only provides effective regularization but also significantly improves performance on smaller cross-family datasets, enabling competitive results. Our model achieves state-of-the-art performance, highlighting the importance of incorporating biologically informed priors into deep learning frameworks for accurate RNA secondary structure prediction.

## References

[1] Stefanie A Mortimer, Mary Anne Kidwell, and Jennifer A Doudna. Insights into rna structure and function from genome-wide studies. Nature Reviews Genetics, 15(7):469–479, 2014.

[2] Kalli Kappel, Kaiming Zhang, Zhaoming Su, Andrew M Watkins, Wipapat Kladwang, Shanshan Li, Grigore Pintilie, Ved V Topkar, Ramya Rangan, Ivan N Zheludev, et al. Accelerated cryo-em-guided determination of three-dimensional rna-only structures. Nature methods, 17(7):699–707, 2020.

[3] Kengo Sato, Manato Akiyama, and Yasubumi Sakakibara. Rna secondary structure prediction using deep learning with thermodynamic integration. Nature communications, 12(1):941, 2021.

[4] David H Mathews, Matthew D Disney, Jessica L Childs, Susan J Schroeder, Michael Zuker, and Douglas H Turner. Incorporating chemical modification constraints into a dynamic programming algorithm for prediction of rna secondary structure. Proceedings of the National Academy of Sciences, 101(19):7287–7292, 2004.

[5] Xinshi Chen, Yu Li, Ramzan Umarov, Xin Gao, and Le Song. Rna secondary structure prediction by learning unrolled algorithms. arXiv preprint 2002.05810, 2020.

[6] Laiyi Fu, Yingxin Cao, Jie Wu, Qinke Peng, Qing Nie, and Xiaohui Xie. Ufold: fast and accurate rna secondary structure prediction with deep learning. Nucleic acids research, 50(3):e14–e14, 2022.

[7] Weijie Yin, Zhaoyu Zhang, Liang He, Rui Jiang, Shuo Zhang, Gan Liu, Xuegong Zhang, Tao Qin, and Zhen Xie. Ernie-rna: An rna language model with structure-enhanced representations. bioRxiv, pages 2024–03, 2024.

[8] Ning Wang, Jiang Bian, Yuchen Li, Xuhong Li, Shahid Mumtaz, Linghe Kong, and Haoyi Xiong. Multi-purpose rna language modelling with motif-aware pretraining and type-guided fine-tuning. Nature Machine Intelligence, pages 1–10, 2024.

[9] Rafael Josip Penić, Tin Vlašić, Roland G Huber, Yue Wan, and Mile Šikić. Rinalmo: General-purpose rna language models can generalize well on structure prediction tasks. arXiv preprint 2403.00043, 2024.

[10] Ziheng Yang. Paml 4: phylogenetic analysis by maximum likelihood. Molecular biology and evolution, 24(8):1586–1591, 2007.

[11] John Jumper, Richard Evans, Alexander Pritzel, Tim Green, Michael Figurnov, Olaf Ronneberger, Kathryn Tunyasuvunakool, Russ Bates, Augustin Žídek, Anna Potapenko, et al. Highly accurate protein structure prediction with alphafold. nature, 596(7873):583–589, 2021.

[12] Josh Abramson, Jonas Adler, Jack Dunger, Richard Evans, Tim Green, Alexander Pritzel, Olaf Ronneberger, Lindsay Willmore, Andrew J Ballard, Joshua Bambrick, et al. Accurate structure prediction of biomolecular interactions with alphafold 3. Nature, pages 1–3, 2024.

[13] Shay Zakov, Yoav Goldberg, Michael Elhadad, and Michal Ziv-Ukelson. Rich parameterization improves rna structure prediction. Journal of Computational Biology, 18(11):1525–1542, 2011.

[14] Zhen Tan, Yinghan Fu, Gaurav Sharma, and David H Mathews. Turbofold ii: Rna structural alignment and secondary structure prediction informed by multiple homologs. Nucleic acids research, 45(20):11570–11581, 2017.

[15] Mehdi Saman Booy, Alexander Ilin, and Pekka Orponen. RNA secondary structure prediction with convolutional neural networks. BMC Bioinformatics, 23(1):58, February 2022.

[16] Padideh Danaee, Mason Rouches, Michelle Wiley, Dezhong Deng, Liang Huang, and David Hendrix. bprna: large-scale automated annotation and analysis of rna secondary structure. Nucleic acids research, 46(11):5381–5394, 2018.

[17] Peter W Rose, Andreas Prlić, Ali Altunkaya, Chunxiao Bi, Anthony R Bradley, Cole H Christie, Luigi Di Costanzo, Jose M Duarte, Shuchismita Dutta, Zukang Feng, et al. The rcsb protein data bank: integrative view of protein, gene and 3d structural information. Nucleic acids research, page gkw1000, 2016.

[18] Ioanna Kalvari, Eric P Nawrocki, Nancy Ontiveros-Palacios, Joanna Argasinska, Kevin Lamkiewicz, Manja Marz, Sam Griffiths-Jones, Claire Toffano-Nioche, Daniel Gautheret, Zasha Weinberg, et al. Rfam 14: expanded coverage of metagenomic, viral and microrna families. Nucleic Acids Research, 49(D1):D192–D200, 2021.

